# An optimized workflow for single-cell transcriptomics and repertoire profiling of purified lymphocytes from clinical samples

**DOI:** 10.1101/803031

**Authors:** Richa Hanamsagar, Timothy Reizis, Mathew Chamberlain, Robert Marcus, Frank O. Nestle, Emanuele de Rinaldis, Virginia Savova

**Affiliations:** Sanofi Immunology and Inflammation Research Therapeutic Area, 270 Albany St, Cambridge, MA 02139; New York University College of Arts and Sciences, 32 Waverly Pl, New York, NY 10003

**Author notes:** Corresponding Author: Virginia Savova, Ph. D., Sanofi, Principal Senior Scientist/Lab Head, Precision Immunology, Immunology & Inflammation Research Therapeutic Area. **Abbreviations: SC**: Single-cell; **PBMCs**: Peripheral Blood Mononuclear Cells; **GEX**: Gene expression; **TB**: Trypan Blue; **AO/PI**: Acridine Orange/Propidium Iodide; **RBCs**: Red Blood Cells; **FSC**: Forward Scatter; **SSC**: Side Scatter.

## Abstract

Establishing clinically relevant single-cell (SC) transcriptomic workflows from cryopreserved tissue is essential to move this emerging immune monitoring technology from the bench to the bedside. Improper sample preparation leads to detrimental cascades, resulting in loss of precious time, money and finally compromised data. There is an urgent need to establish protocols specifically designed to overcome the inevitable variations in sample quality resulting from uncontrollable factors in a clinical setting. Here, we explore sample preparation techniques relevant to a range of clinically relevant scenarios, where SC gene expression and repertoire analysis are applied to a cryopreserved sample derived from a small amount of blood, with unknown or partially known preservation history. We compare a total of ten cell-counting, viability-improvement, and lymphocyte-enrichment methods to highlight a number of unexpected findings. Trypan blue-based automated counters, typically recommended for single-cell sample quantitation, consistently overestimate viability. Advanced sample clean-up procedures significantly impact total cell yield, while only modestly increasing viability. Finally, while pre-enrichment of B cells from whole peripheral blood mononuclear cells (PBMCs) results in the most reliable BCR repertoire data, comparable T-cell enrichment strategies distort the ratio of CD4+ and CD8+ cells. Furthermore, we provide high-resolution analysis of gene expression and clonotype repertoire of different B cell subtypes. Together these observations provide both qualitative and quantitative sample preparation guidelines that increase the chances of obtaining high-quality single-cell transcriptomic and repertoire data from human PBMCs in a variety of clinical settings.

## Introduction

Single-cell analysis has become increasingly popular in the field of cancer immunology [1] and autoimmune disorders [2, 3], with the aim to potentially identify patient-specific signatures and apply a more targeted therapy [4–6]. There is also enhanced focus on T and B lymphocyte profiling in infections [7], or in patients treated with vaccines [8] or antibody-based immunotherapies [9]. Additionally, studies have also investigated antibody repertoires in patients with autoimmune disorders [10, 11]. Inspired by these early efforts, large disease-focused consortia are increasingly investing in SC transcriptomics on human biological samples due to the broad readout that this technology can provide using only a small amount of tissue as input (cf. Accelerating Medicines Partnership (AMP), Open Targets) [12].

While the potential of single-cell approaches for bench-to-bedside is evident, its future applicability depends to a large extent on the successful development of robust sample preparation techniques [13]. There is an emergent need to establish protocols and workflows optimized for clinical settings. These must be specifically designed to overcome the inevitable variations in sample quality resulting from uncontrollable factors in sample collection and preservation. Failure to account for this can result in low quality data, as well as loss of precious samples, time and ultimately money.

SC transcriptomics is a time-consuming and expensive procedure from sample collection, to single cell encapsulation, to library preparation and finally sequencing and downstream data analysis. The first step of this workflow -- sample preparation, is of utmost importance. Sample collection techniques in a clinical setting typically vary from person-to-person, from site-to-site and from day-to-day. Adoption of standardized protocols for sample preparation are required to address technical variability and accurate sample clean-up steps are required to increase viability and preserve phenotype post-thawing.

Here, we explore and compare sample preparation techniques applicable to a range of clinically relevant hypothetical scenarios, where the starting point is a PBMC sample derived from anywhere from 20ml to 0.5 ml of blood, with unknown or partially known preservation history (Fig. 1). The number of cells and viability of such a sample cannot be well-established in advance, and once thawed, the sample can be profiled using three different 10X Genomics® single-cell workflows: whole PBMC 5’ gene expression (GEX) analysis, and/or enriched B/T-cell profiling using GEX and/or V(D)J kits, depending on research need. This scenario raises a number of questions in sample preparation, specifically: how to accurately estimate cell count and viability; how to improve viability; whether and how to enrich B and T cells.

**Figure 1.**
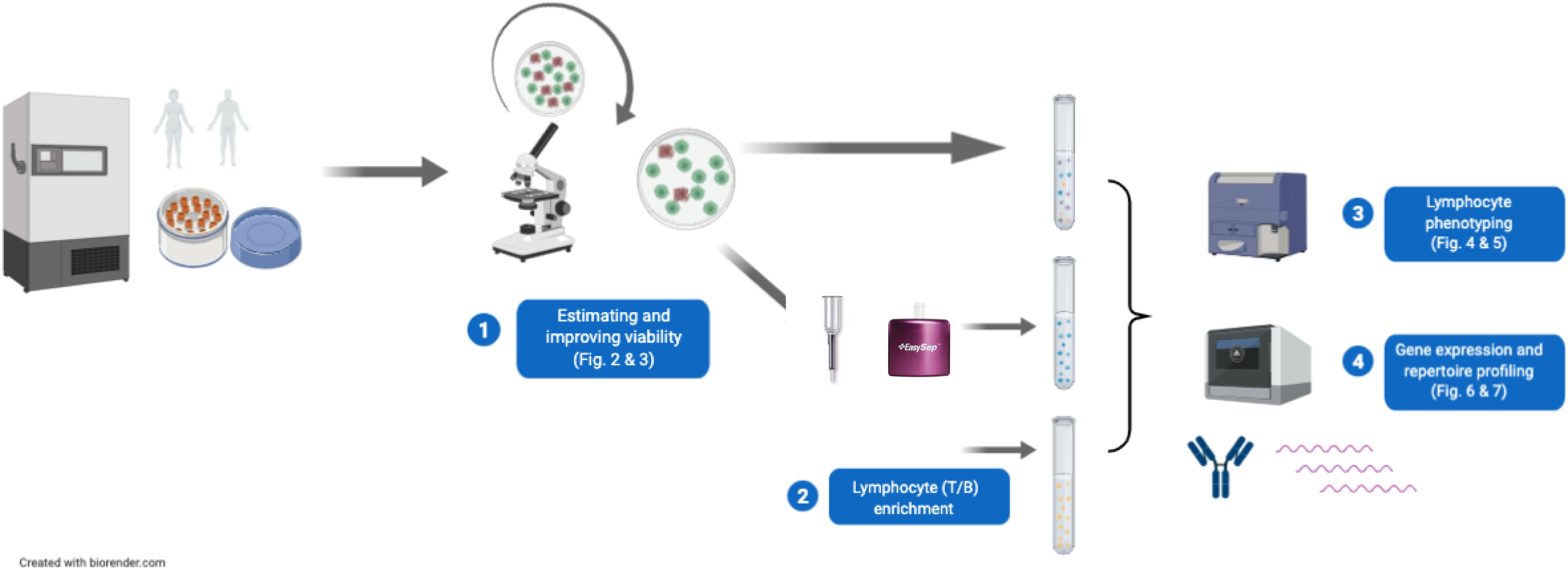
Optimizing single-cell sample preparation and workflow using previously frozen human PBMCs. First, we tested different methods for estimating and improving viability on freshly thawed human PBMCs. The results for these experiments are summarized in Figures 2 and 3. Next, an aliquot of whole PBMCs was set aside. Another aliquot of PBMCs was subjected to different methods of T and B lymphocyte enrichment. Following this, the aliquot of whole PBMCs, as well as the enriched cells were stained with fluorescence antibodies and the yield and purity of enriched cells was assessed by flow cytometry. These results are summarized in Figures 4 and 7. Finally, 10X Genomics® single-cell 5’ gene expression and V(D)J repertoire profiling was performed in order to assess the differences in gene expression/repertoire data obtained from whole PBMCs vs. enriched lymphocytes, results for which are summarized in Figures 5 and 6.

We found that manual cell counting using hemocytometer and trypan blue is the most accurate way to count cell and obtain viability data. Fluorescence-based automated counters perform similarly to hemocytometer-based manual counting, but trypan blue-based automated counters, on which 10X Genomics® sample recommendations are based, consistently overestimate viability. Furthermore, sample clean-up procedures significantly impact total cell yield while increasing viability of cell suspension, hence care should be taken while working with precious samples. Finally, pre-enrichment of B cells from whole PBMCs results in the most reliable repertoire data as compared with VDJ target-enrichment from whole PBMCs, but comparable T-cell enrichment come with a significant caveat. Overall, we provide a sample preparation guideline for researchers that will increase the chances of obtaining high-quality single-cell transcriptomic and BCR repertoire data from human PBMCs.

## Methods and Materials

### Human Peripheral Blood Mononuclear cell (PBMC) Isolation and Storage

Whole blood (containing sodium citrate as an anti-coagulant) was purchased from MedRACS Clinic Research, (Asentral IRB study no. 2014-327A). ACCUSPIN System-HISTOPAQUE-1077 tubes (Sigma Aldrich, Catalog # A6929) were brought to room temperature (RT) and centrifuged at 800 g for 30 seconds. Following this, 35 ml of whole blood was layered on top of the frit in each Accuspin column and centrifuged at 800 g for 20 min without break. Mononuclear cell layer was formed under the plasma and above the frit. The cells were collected in a sterile 50 ml conical tube using sterile transfer pipette. Cells were washed by filling the tubes to 50 ml with wash buffer (0.5% BSA, 2mM EDTA in 1X PBS) and centrifuged at 300 g for 10 min. Supernatant was aspirated and pellet was gently resuspended with 10 ml wash buffer. The tube was once again topped with wash buffer up to 50 ml mark and centrifuged at 300 g for 10 min. The wash was repeated one more time. After this, pellet was resuspended in 10 ml wash buffer and pooled (if multiple sample tubes were present per donor). Cell count and viability was determined using Vi-Cell XR Viability Analyzer. Cells were pelleted by centrifuging at 300 g for 10 min. Supernatant was aspirated and resuspended in FBS containing 10% DMSO at a concentration of 20 million per ml. Cell suspension was transferred to cryotubes (2 ml per tube) and placed in Mr. Frosty Cryo 1 °C freezing container (ThermoScientific, Cat# 5100-001) and stored at −80 °C for 24 hours. Following this, tubes were transferred to liquid nitrogen tank for long-term storage.

### Cell Counting Optimization

#### Single Cell Suspension

A total of 4 different frozen, healthy human PBMCs samples were analyzed in this experiment. Frozen vials containing human PBMCs were thawed for 2 min in water bath at 37 C. After this, cell suspension was transferred to a fresh 2 ml Eppendorf tube using wide bore pipette tip (Thermo Scientific FINNTIP). Sample was centrifuged (Eppendorf 5417R) at 300 g for 5 min at 4 C. Supernatant was removed, and 2 ml of 0.04% BSA/PBS (Bovine Serum Albumin – Fisher BioReagents Cat# BP9703-100; 1X DPBS – GIBCO Cat# 14190-136) was added. Pellet was gently resuspended using wide-bore pipette tip and re-centrifuged at 300g, 5 min, 4 C. Supernatant was removed, and cell pellet was resuspended in 1 ml of 0.04% BSA/PBS using wide-bore pipette tip. Serial dilutions of the cell suspension were made as follows: 1:1, 1:2, 1:4, 1:8, 1:16 and 1:32 using 0.04% BSA/PBS, and placed on ice.

#### Vi-cell XR

100 ul of cell suspension for each of the dilutions including undiluted (1:0) was mixed with 400 ul of 1X PBS in the ViCell cuvette (Beckman Coulter, Part # 723908). Cell counting was performed based on manufacturer’s instructions (Beckman Coulter Vi-cell XR – 490992, on Vi-CELL XR 2.04.004 Software). Every sample and its dilution were separately counted 3-4 times to get technical replicates. Total number of cells, number of live cells and number of dead cells were noted.

#### BioRad TC20 Automated Cell Counter

Cell suspensions were diluted with trypan blue (Gibco Trypan Blue Stain (0.4%) Cat# 15250 - 061) either 1:10 (10 ul of cells + 90 ul of trypan blue, dilution factor or DF = 10) or 1:1 (10 ul of cells + 10 ul of of trypan blue, DF = 2), and 10 ul of mixture was pipetted into the chamber of BIO RAD Counting Slides (BioRad, Cat# 145-0011). Slide was inserted into the instrument (BIO RAD TC20 Automated Cell Counter) and counts were noted down. Cell counts were further multiplied by the dilution factor to get final cell count/ml. Every sample and its dilution were separately counted 3-4 times to get technical replicates. Total number of cells, number of live cells and number of dead cells were noted.

#### Manual Hemacytometer-based cell counting

Cell suspensions were diluted with trypan blue (Gibco Trypan Blue Stain (0.4%) REF 15250 - 061) either 1:10 (10 ul of cells + 90 ul of trypan blue, dilution factor = 10) or 1:1 (10 ul of cells + 10 ul of of trypan blue, dilution factor = 2), and 10 ul of mixture was pipetted into the chamber of in CYTO C-Chip (inCYTO C-Chip, DHC-N01-5, Neubauer Improved). Cells were counted using a brightfield microscope at 20X magnification. Cells that appeared blue in color were counted as dead cells, whereas cells that were transparent were counted as live cells. Cells were also counted on the edges of the grids, and careful measures were taken to avoid counting debris or red blood cells. Cells were counted in 4 corner quadrants, averaged, and multiplied by 10,000 times dilution factor to get cells/ml count. Two separate researchers counted each slide separately, and cell numbers were averaged across researchers. Equation for determining cell number per ml is as follows:

(Q1+Q2+Q3+Q4)/4 * 10,000 * Dilution Factor = number of cells per ml Every sample and its dilution were separately counted 3-4 times by 2 separate researchers to get technical replicates. Total number of cells, number of live cells and number of dead cells were noted.

#### Cellaca MX

Cellaca MX Automated Cell Counter (Nexcelom Bioscience) was used as per manufacturer’s instructions. Briefly, cells were diluted with Cellometer ViaStain^TM^ AOPI Staining Solution (Catalog # CS2-0106-5ML) in a dilution of either 1:1 or 1:10 and then plated into the counting wells of the 24-well Counting plate (Product # CHM24-A100-001). Plate was loaded into the Cellaca MX instrument and focus was optimized. Every sample and its dilution were separately counted 3-4 times to get technical replicates. Total number of cells, number of live cells and number of dead cells were noted.

#### LUNA-FL™ Dual Fluorescence Cell Counter

LUNA-FL™ Dual Fluorescence Cell Counter (Cat # L20001, Logos Biosystems) was used as per manufacturer’s instructions. Briefly, cells were diluted with Acridine Orange/Propidium Iodide Stain (Cat # F23001, Logos Biosystems) in a dilution of either 1:1 or 1:10 and then pipetted onto LUNA™ Cell Counting Slides (Cat # L12001, Logos Biosystems). Slide was loaded onto the instrument and focus was optimized using the focusing knob on the side. Every sample and its dilution were separately counted 3-4 times to get technical replicates. Total number of cells, number of live cells and number of dead cells were noted.

### Sample Clean-up Optimization

#### Single Cell Suspension Preparation

A total of 4 different frozen human PBMCs samples were analyzed in this experiment. Frozen vials containing cells were thawed for 2 min in water bath at 37C. After this, cell suspension was transferred to a fresh 2 ml Eppendorf tube using wide bore pipette tip (Thermo Scientific FINNTIP). Sample was centrifuged (Eppendorf 5417R) at 300 g for 5 min at 4C. Supernatant was removed, and 2 ml of 0.04% BSA/PBS (BSA – Fisher BioReagents BP9703-100) (PBS – Gibco DPBS 1X REF. 14190-136) was added. Pellet was gently resuspended using wide-bore pipette tip. Total number of cells, number of live and dead cells were counted using manual hemacytometer. The sample was split into 4 tubes of 500 μl each. Each of the tubes was used for 1 method of sample clean-up.

#### Dead Cell removal methods

##### Method 1: MACS Miltenyi Biotec Dead Cell Removal Kit (Cat # 130-090-101)

Manufacturer’s instructions were followed for removing dead cells. Briefly, cells were centrifuged at 300 g for 5 minutes. Supernatant was completely removed, and cell pellet was resuspended in 100 ul of Dead Cell Removal MicroBeads per 10 cells, mixed and incubated at room temperature for 15 min. MS columns (Cat # 130-042-201) were attached to OctoMACS magnetic separator (Cat # 130-042-109) and primed by rinsing with 500 ul of 1X Binding buffer. Cell suspension was applied to the column and washed 4 times with 500 ul of 1X Binding buffer. Effluent containing primarily live cells was collected, centrifuged at 300 g for 5 min, resuspended in cold 0.04% BSA/PBS and counted manually using hemacytometer.

##### Method 2: StemCell Technologies EasySep Dead Cell Removal (Annexin V) Kit (Catalog #17899)

Manufacturer’s instructions were followed for removing dead cells. Briefly, cells were centrifuged at 300 g for 5 minutes. Supernatant was completely removed and resuspended in 1X PBS containing 2% fetal bovine serum (FBS) and 1 mM CaCl_2_ at a concentration of 10 per ml. Sample was transferred into a 5 ml polystyrene tube. EasySep Dead Cell Removal (Annexin V) Cocktail (Cat. 17899C) and EasySep Biotin Selection Cocktail (Cat. 18153) were added, mixed and incubated at room temperature for 3 min. EasySep Dextran RapidSpheres (Cat. 50103) were vortexed and added, after which final volume was made up to 2.5ml using 1X PBS/FBS/CaCl_2_ solution above. Tube was placed on the EasySep Magnet (Cat. 18000) for 3 min and cell suspension was carefully decanted into a new tube. Decanted solution primarily containing live cells was centrifuged at 300 g for 5 min, resuspended in cold 0.04% BSA/PBS and counted manually using hemacytometer.

##### Method 3: MACS Miltenyi Biotec Debris Removal Solution (Cat. 130-109-398)

Manufacturer’s instructions were followed for removing debris from cells. Briefly, cells were centrifuged at 300 g for 10 minutes at 4 C. Supernatant was completely removed, cell pellet was resuspended in 1 ml of cold 1X PBS, 300 uL of Debris Removal Solution was added, transferred to a 15 ml tube and mixed well. The solution was gently overlayed with 1 ml of cold 1X PBS. Sample was centrifuged at 4 C, 300 g for 10 min with full acceleration and full brake. Top two layers were aspirated and discarded. The bottom layer was left undisturbed, and volume was made up to 15 ml with cold 1X PBS. Cells were mixed gently and centrifuged at 1000g, for 10 min at 4C. Supernatant was removed, and cells were resuspended in cold 0.04% BSA/PBS for counting manually using hemacytometer.

##### Method 4

10X Genomics® recommended washes: Cells were washed 3 times with 0.04% BSA/PBS at 150 g, 5 min, 4 C. After the final spin, supernatant was completely removed, and cell pellet was resuspended in 1 ml of cold 0.04% BSA/PBS for counting manually using hemacytometer.

### T and B cell Enrichment Optimization

#### Cell preparation

A total of 4 different frozen human PBMCs samples were analyzed in this experiment. Frozen vials containing cells were thawed for 2 min in water bath at 37 C. After this, cell suspension was transferred to a fresh 2 ml Eppendorf tube using wide bore pipette tip (Thermo Scientific FINNTIP). Sample was centrifuged (Eppendorf 5417R) at 300 g for 5 min at 4 C. Supernatant was removed, and 2 ml of 0.04% BSA/PBS was added. Pellet was gently resuspended using wide-bore pipette tip and the washes were repeated for additional 2 times (total of 3 washes). Cells were counted manually using hemacytometer. A small aliquot of cells was set aside for direct staining and analysis of whole PBMCs by flow. The rest of the cells were equally divided into two volumes, one for MACS Miltenyi Biotec enrichment and other for STEMCELL enrichment kit.

#### B cell enrichment

##### MACS Miltenyi Biotec Pan B Cell Isolation Kit, human (Cat# 130-101-638)

Protocol was followed as per manufacturer’s instructions. Briefly, PBMCs were counted, centrifuged at 300 g for 5 min at 4 C, and resuspended in 40 ul of MACS buffer [prepared by diluting MACS® BSA Stock Solution (# 130-091-376) 1:20 with autoMACS® Rinsing Solution (# 130-091-222)] per 10 cells. Then, 10 ul of Pan B cell Biotin-antibody cocktail was added per 10 cells. Cells were mixed and incubated for 5 min in the fridge. Following this, 30 ul of MACS buffer and 20 ul of anti-biotin Microbeads were added. Cells were mixed and incubated in the fridge for 10 min. MS columns (Cat # 130-042-201) were used for separation. Columns were placed on the OctoMACS magnetic separator (Cat # 130-042-109) and prepared by rinsing with 500 μl of MACS buffer. Cell suspension was applied to the column and all solution was allowed to flow through. Columns were further washed one with 500 ul MACS buffer The eluent containing enriched B cells were collected and centrifuged at 300 g for 5 min at 4 C. Pellet was resuspended in 1 ml of 1X PBS buffer for counting and flow cytometry.

##### STEMCELL^TM^ Technologies EasySep™ Human B Cell Enrichment (Cat #17954)

Protocol was followed as per manufacturer’s instructions. Briefly, cells were counted, centrifuged at 300 g for 5 min at 4° C. Pellet was resuspended in EasySep buffer (Catalog #20144) and incubated with DNase I solution (Catalog #07900) at a concentration of 100 ug/ml at room temperature for 15 min prior to labelling and separation. Following incubation, any cell aggregates were filtered out using 37 um cell strainer (Catalog #27250). Cells were then diluted to a concentration of 5×10^7^ cells/ml in EasySep buffer. Sample was transferred to 5 ml polystyrene round-bottom tube (Catalog #38007) and enrichment cocktail was added at a concentration of 50 ul/ml. Sample was mixed and incubated at room temperature for 10 min. Magnetic particles were thoroughly vortexed and added to the sample at a concentration of 75 ul/ml and incubated at room temperature for 5 min. Following this, the cell solution was topped up to 2.5 ml with EasySep buffer, mixed and placed on the EasySep magnet (Catalog #18000) for 5 min at room temperature. Enriched B cells were collected by pouring off in one quick motion and centrifuged and centrifuged at 300 g for 5 min at 4 C. Pellet was resuspended in 1 ml of 1X PBS buffer for counting and flow cytometry.

#### T cell enrichment

##### MACS Miltenyi Biotec Pan T Cell Isolation Kit, human (Cat# 130-096-535)

Protocol was followed as per manufacturer’s instructions. Briefly, PBMCs were counted, centrifuged at 300 g for 5 min at 4° C, and resuspended in 40 ul of MACS buffer [prepared by diluting MACS® BSA Stock Solution (# 130-091-376) 1:20 with autoMACS® Rinsing Solution (# 130-091-222)] per 10^7^ cells. Then, 10 ul of Pan T cell Biotin-antibody cocktail was added per 10^7^ cells. Cells were mixed and incubated for 5 min in the fridge. Following this, 30 ul of MACS buffer and 20 ul of anti-biotin Microbeads were added. Cells were mixed and incubated in the fridge for 10 min. MS columns (Cat # 130-042-201) were used for separation. Columns were placed on the OctoMACS magnetic separator (Cat # 130-042-109) and prepared by rinsing with 500 ul of MACS buffer. Cell suspension was applied to the column and all solution was allowed to flow through. Columns were further washed one with 500 ul MACS buffer. The eluent containing enriched T cells were collected and centrifuged at 300 g for 5 min at 4 C. Pellet was resuspended in 1 ml of 1X PBS buffer for counting and flow cytometry.

##### STEMCELL™ Technologies EasySep™ Human T Cell Isolation Kit (Cat #17951)

Protocol was followed as per manufacturer’s instructions. Briefly, cells were counted, centrifuged at 300 g for 5 min at 4°C. Pellet was resuspended in EasySep buffer (Catalog #20144) and incubated with DNase I solution (Catalog #07900) at a concentration of 100 ug/ml at room temperature for 15 min prior to labelling and separation. Following incubation, any cell aggregates were filtered out using 37 μm cell strainer (Catalog #27250). Cells were then diluted to a cencentration of 5×10^7^ cells/ml in EasySep buffer. Sample was transferred to 5 ml polystyrene round-bottom tube (Catalog #38007) and isolation cocktail was added at a concentration of 50 μl/ml. Sample was mixed and incubated at room temperature for 5 min. RapidSpheres^TM^ were thoroughly vortexed and added to the sample at a concentration of 40 μl/ml. Following this, the cell solution was topped up to 2.5 ml with EasySep buffer, mixed and placed on the EasySep magnet (Catalog #18000) for 3 min at room temperature. Isolated T cells were collected by pouring off in one quick motion and centrifuged at 300 g for 5 min at 4 C. Pellet was resuspended in 1 ml of 1X PBS buffer for counting and flow cytometry.

##### Flow Cytometry

Bio-Rad Cell Analyzer (ZE5, 5 lasers, Cat # 12004279) was used for all flow cytometry experiments. Compensation was done prior to assay setup on the flow cytometer. Briefly, 2 drops of the Ultracomp beads (Invitrogen, Cat# 01-2222-42) were added to a 5ml polystyrene Falcon round bottom tube, one tube per stain. An unstained tube was prepared with just 2 drops of beads and 400ul of stain buffer (2% FBS in 1XPBS). Single color tubes were vortexed and incubated on ice for 15-20 minutes in the dark. After incubation, 300ul of stain buffer was added per tube and vortexed to mix. Unstained and single-color beads were then used to set appropriate voltages on the Bio-Rad ZE5. Cells were centrifuged at 300g, 12 C for 5 minutes. Supernatants were removed and 100ul of stain panel was added to each cell pellet, resuspended and allowed to incubate on ice, in the dark for 30 minutes. Following this, cells were centrifuged as above, supernatant was discarded, and cell pellet was resuspended in 200 ul of 7-AAD Viability stain (5 ul 7AAD + 195 ul staining buffer), and incubated on ice for 10 min in the dark. After the incubation with viability stain, samples are ready for acquisition on the cytometer. All flow analysis was performed on FlowJo v10.

For more information on the antibodies used and representative gating strategy, please refer to Supplementary Figure 1. For detailed information on flow analysis please refer to Supplementary Information.

### Single-Cell workflow

#### Cell preparation

Frozen vials containing cells were thawed for 2 min in water bath at 37 C. After this, cell suspension was transferred to a fresh 2 ml Eppendorf tube using wide bore pipette tip (Thermo Scientific FINNTIP). Sample was centrifuged (Eppendorf 5417R) at 300 g for 5 min at 4 C. Supernatant was removed, and 2 ml of 0.04% BSA/PBS was added. Pellet was gently resuspended using wide-bore pipette tip and the washes were repeated for additional 2 times (total of 3 washes). Cells were counted manually using hemacytometer. Based on the downstream processing, either cells were directly diluted and processed through 10X Genomics® workflow or further enriched for T and B cells and then processed through 10X Genomics® workflow.

##### 10X Genomics®: 5’ Gene expression and VDJ repertoire immune profiling

Samples were processed as per Chromium Single Cell V(D)J Reagent Kits Protocol by 10X Genomics®. Approximately 4000 cells were targeted for each sample recovery. For whole PBMCs, a part of cDNA was set aside for BCR target enrichment and VDJ repertoire library preparation. Another part of cDNA from PBMCs was processed for 5’ gene expression library. The quality and quantity cDNA and end libraries was assessed on the 2100 Bioanalyzer instrument (Agilent) using high sensitivity DNA Chips and Kit (Agilent, Catalog # 5067-4626). For a more accurate quantification of libraries, qPCR was performed after serial dilution of the libraries 1:10,000, 1:20,000, 1:40,000 and 1:80,000 in 10 mM Tris-HCl. Kapa Library quantification kit Illumina® Platforms (Kapa Biosystems, Catalog # KK4824 – 07960140001) was used for qPCR on QuantStudio 7 Flex Real-Time PCR system. The qPCR cycling protocol was as per Kapa Library quantification kit protocol [14]. Libraries were pooled in an equimolar fashion to obtain a final library pool of 10nM. Gene expression libraries and V(D)J repertoire libraries were pooled separately as they require different sequencing metrics.

Sequencing was performed on NOVAseq 6000 system (Illumina®), with NovaSeq 6000 S1 Reagent Kit (300 cycles) (Catalog # 20012863). Sequencing depth and cycle number was as per 10X genomics recommendations. Specifically, for V(D)J enriched libraries, read 1 = 150 cycles, i7 index = 8 cycles and read 2 = 150 cycles. For 5’ gene expression libraries, read 1 = 26 cycles, i7 index = 8 cycles and read 2 = 98 cycles. For V(D)J enriched libraries, we performed a sequencing depth of 5000 read pairs per cell and for 5’ gene expression libraries we did 25,000 read pairs per cell.

#### Single Cell Data analysis

Following BCL conversion, fastq files were processed through CellRanger Pipeline (10X Genomics®). This allowed demultiplexing, alignment, filtering, barcode counting, and UMI counting, and generating of cell x barcode matrices. These matrices were then input into SPRING [15], which is a tool for visualizing and analyzing single-cell data [16]. Using this, we were able to visualize single-cell data obtained from whole PBMCs and enriched B cells. Using the gene-finder tool, cells expressing gene of interested were highlighted and compared across groups. Differential gene expression analysis (DEG) was performed with the R package Seurat [17] (v3.0.0) with the FindMarkers function using the default settings.

#### Data analysis and Statistics

FlowJo (version 10) was used for analyzing flow data. GraphPad Prism (version 8) was used for generating graphs and performing statistical analyses. For the sake of organization, all statistical analyses and methods are detailed in supplementary worksheet “Statistical Analysis”. Levels of significance are indicated by: ***p<0.001, **p<0.01 and *p<0.05. For differential gene expression analysis, statistical tests are summarized in worksheet “Supplementary_DEG”.

## Results

### Trypan Blue-based manual hemacytometer provides the most accurate cell numbers and viability data

Often times, clinical samples are stored at less than ideal conditions, for long periods of time, and hence might be of poor quality at the time of processing. In addition, clinical samples can be precious, hence it is important to extract maximum amount of data from a small amount of sample. Low viability is one of the most common challenges faced while handling clinical samples. The Single Cell Protocols Cell Preparation Guide by 10X Genomics® [18] and inDrop^TM^ Single Cell Encapsulation and RT protocol (Version 2.4) [19] by 1CellBio recommends a sample viability of 90% and 95%, respectively. However, the viability reading can vary largely based on the method used for counting cells. 10X Genomics® recommends the use of Countess® II Automated Cell Counter for most applications. In the absence of an automated method, 10X Genomics® recommends using manual hemacytometer, which is also the method recommended by 1CellBio.

To test the best possible method for accurately counting human PBMCs, we first compared three trypan-blue (TB) based counting methods: (1) hemacytometer based manual counting, (2) Vi-CellXR Viability Analyzer and (3) BioRad TC20 Automated Cell Counter. Two-way ANOVA revelated that TB-based automated counters significantly overestimated number of live cells at higher concentrations i.e. at dilutions 0 and 1:1 (Fig 2A), significantly underestimated the number of dead cells at all concentrations, except at dilution of 1:32 (Fig. 2B) and consequently significantly overestimated the viability of cells at all concentrations compared to manual hemacytometer counting (Fig. 2C). There was no significant difference observed in total counts obtained from automated counters vs. manual counting, except at dilution 1:32 (Suppl. Fig. 2A). For 10X Genomics® workflow, it is recommended to have an optimum cell concentration from 0.7 to 1.2 million cells per ml, prior to diluting with reverse transcriptase enzyme mix. This ensures that the actual cell recovery is as close as possible to the target cell recovery. For this reason, we performed additional replicates of PBMCs to fall within the dilution range of 1:2 and 1:4, and that corresponded to approximately 1 million per ml concentration for our particular cell suspension sample. At this concentration range, the TC20 Automated Cell Counter significantly overestimated live cells (Fig. 2A-bottom), both counters underestimated dead cells (Fig. 2B-bottom) and overestimated viability (Fig 2C-bottom). The total cell count per ml was only different at 1:2 dilution between the 2 automated methods (Suppl. Fig. 2B). But no difference was seen at any dilution when compared with manual hemacytometer counts. Next we compared cell counts obtained from TB-based manual counting with that of two automated AO/PI-based fluorescence counters: (1) Cellaca MX and (2) Luna-FL. Interestingly, there was no significant difference in dead cells/ml (Fig. 2E) between the AO/PI based automated counters and manual counting. But for live cells per ml (Fig. 2D), there was an overall significant difference (*p = 0.037) in “Method of counting” across all dilutions, but no difference in Tukey’s multiple comparison test at any of the individual dilutions (see statistical analysis worksheet). Additionally, there was a significant interaction in % viability values (Fig. 2F, see statistical analysis worksheet), and Tukey’s multiple comparison test revealed a significant difference in % viability between manual counting and Luna-FL at 1:16 dilution.

**Figure 2.**
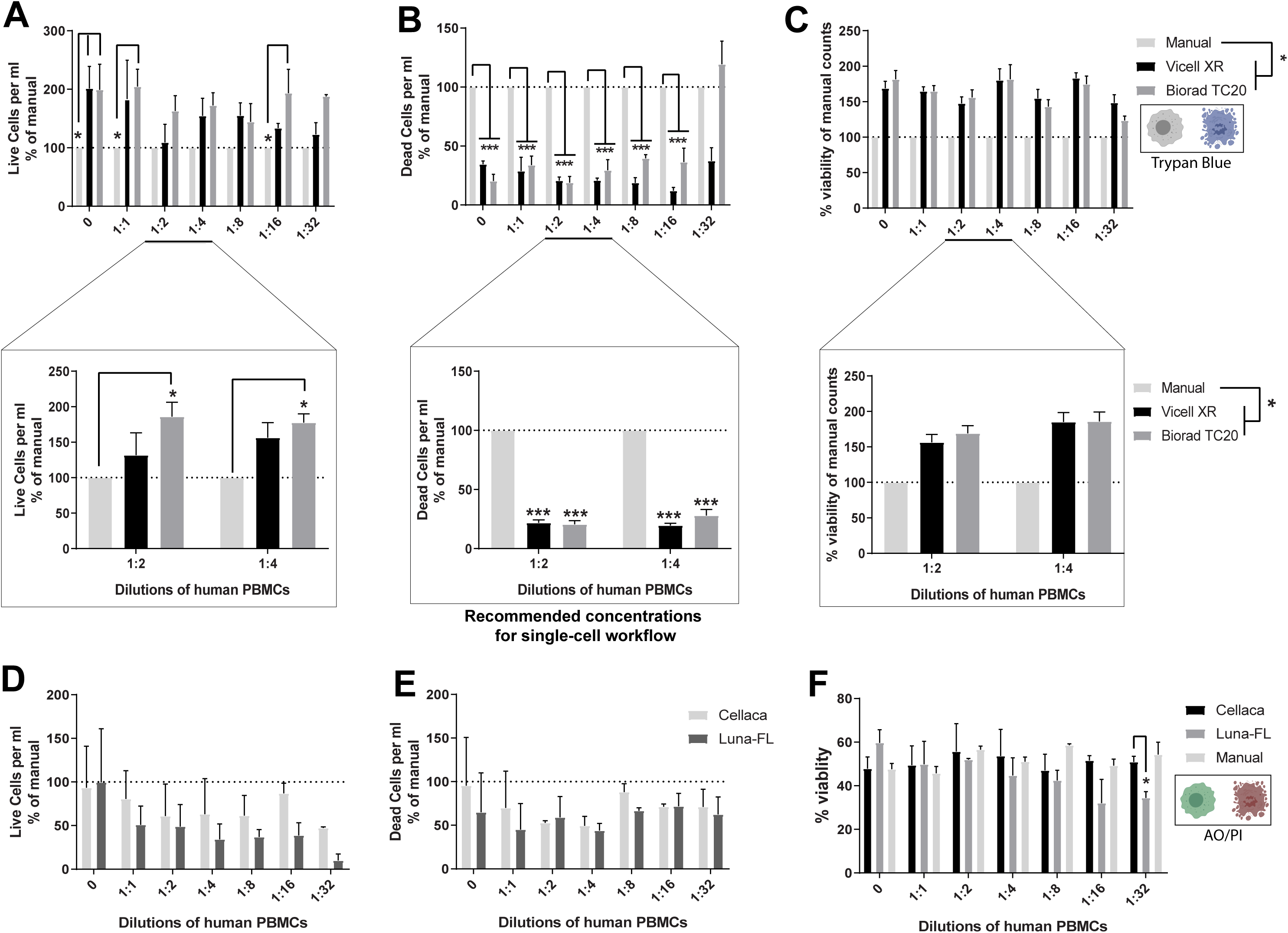
Cell viability estimation using automated TB-based counters or automated AO/PI-based counters. Human PBMCs were diluted using 0.04% BSA/PBS solution and each dilution was counted using Manual hemacytometer, automated TB-based counters or automated AO/PI-based counters. (A-top) Live cells per ml, (B-top) Dead cells per ml, and (C-top) % viability for the TB-based automated methods were noted and calculated as % of manual counting method. Multiple technical replicates were performed for 1:2 and 1:4 dilutions and re-counted as above. (A-bottom) Live cells per ml, (B-bottom) Dead cells per ml, (C-bottom) % viability for the automated counters were reported as % of manual counts. Two-way ANOVA with matched samples across different dilutions, followed by Tukey’s Multiple Comparison test (n=3-6). (D) Live cells per ml and (E) Dead cells per ml were reported at % of manual counts. (F) % viability calculated as live per ml/total per ml*100. Matched sample, Two-way ANOVA followed by Tukey’s multiple comparison test (n=4). For detailed statistics refer to Supplementary information.

### Dead Cell removal steps can result in significant loss in total number of cells

Once an accurate cell count is established, sample clean-up might be necessary in order to increase the viability of the sample. Hence, we tested different dead cell or debris removal methods including: (1) 10X Genomics®-recommended low-spin washes, (2) Magnetic column-based Dead Cell Removal Kit (3) Debris Removal Solution and (4) Column-free Dead Cell Removal Kit. We found that Magnetic-column-based Dead Cell Removal Kit was the most efficient at removing dead cells, by increasing viability significantly compared to no-clean-up control. (Fig.3A, one-way ANOVA), with only slight improvement in viability for all other methods (see statistical analysis worksheet for p-values). All methods resulted in overall cell loss following cleanup, with the most cell loss seen in the two magnetic separation methods, and the least with the washes alone (Fig. 3B, two-way ANOVA).

**Figure 3.**
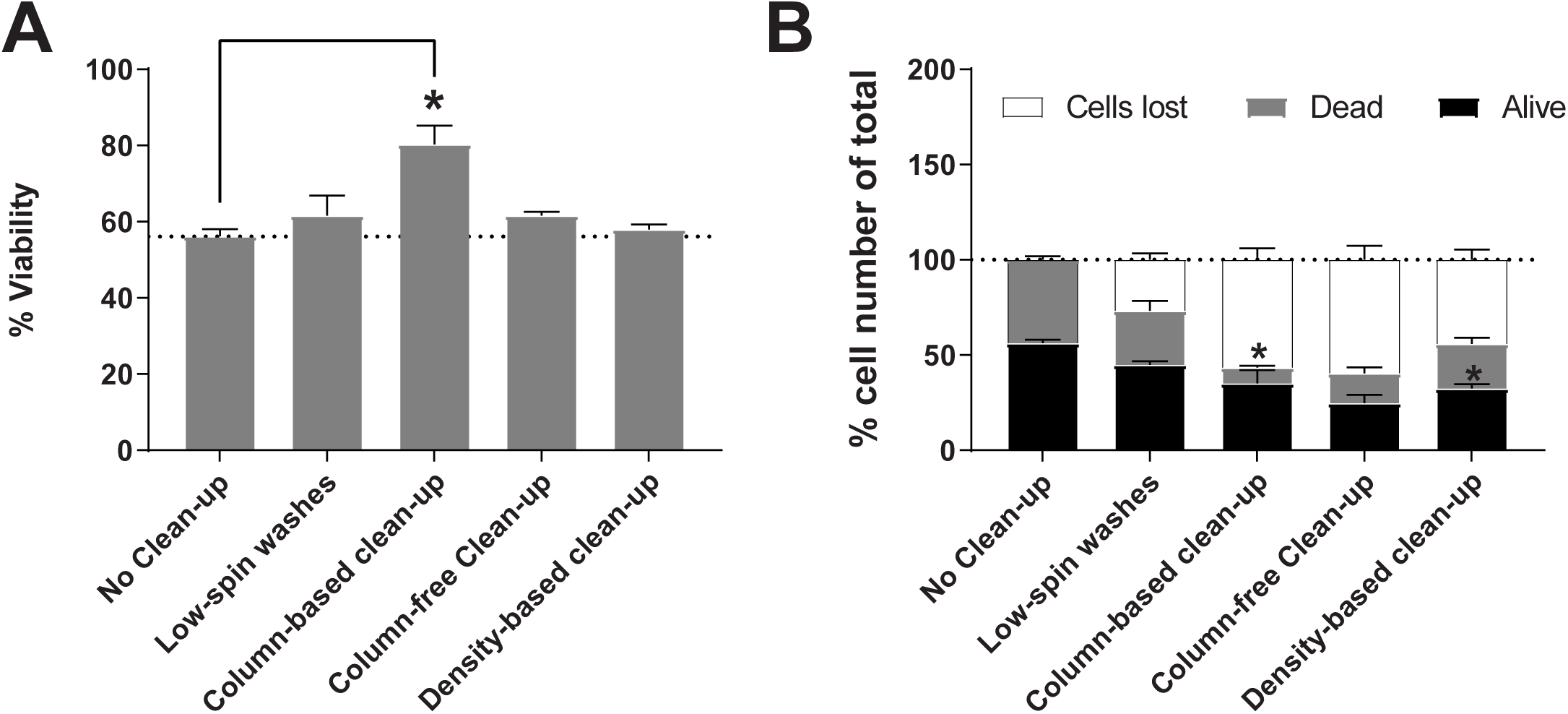
Quantifying cell loss and viability of PBMC samples after applying viability improvement protocols. Human PBMCs were split into 5 equal parts and 4 aliquots were subjected to a different dead cell removal method each. One aliquot was kept aside as “no-cleanup control”. Cell counts were determined using manual hemacytometer before and after dead cell removal. (A) % viability pre and post-clean up (One-way ANOVA). (B) % cells lost, % dead cells and % live cells of total across different methods of clean-up. Two-way ANOVA with matched samples, followed by Tukey’s Multiple Comparison test (n=10). (*p<0.05). For detailed statistics refer to Supplementary information.

### Comparison of performance of pre-enrichment kits based on yield and purity

To test different commercially available kits, T and B cells were enriched from human PBMCs using negative enrichment magnetic separation with column-based or column-free methods. Following this, we stained whole PBMCs and enriched cells with markers of B cells (CD19 and CD20) or T cells (CD3, CD4, CD8) and performed flow analysis to assess purity and yield of enriched cell for each kit (Refer to supplementary Figure 1 for information on gating).

#### T cell enrichment

T cells were negatively enriched from whole PBMCs using either column-based or column-free magnetic separation, stained with antibodies for CD3, CD4 and CD8 and analyzed by flow analysis. First, all cells were gated on FSC-A and FSC-H to remove any doublets. Singlets were further gated on 7-AAD viability stain to exclude any dead cells. Next, live, single cells were gated on CD3 antibody, and only cells positive for CD3 were analyzed further for CD4 and CD8 expression (Fig. 4A). Following flow analysis, we found that in general, both column-free and column-based methods performed comparably in their ability to enrich for T cells (Fig. 4B). As we did not see any bead contamination with the original protocol of column-free T cell isolation method, we did not perform 2 magnetic incubations for this experiment (data not shown). Both the enrichment kits were able to remove contaminating non-T cells from the enriched cells to the same level as shown by a reduction in CD3-cell proportion in the enriched cells (Fig. 4C). Interestingly, however, the ratio of CD4+ to CD8+ cells as measured by FACS was altered in enriched T cells compared to whole PBMCs. Specifically, in whole PBMCs, the ratio of CD4+ to CD8+ cells is less than 1. However, this ratio increased to almost 2 for both the enrichment kits (Fig. 4D). To understand if this was due to difference in relative abundance of CD4+CD8+ double positive cells in whole PBMCs vs. enriched cells, we compared the levels of these cells across all methods. However, there was no difference seen in the proportion of CD4+CD8+ cells in CD3+ cells in the enriched T cells compared to whole PBMCs (Fig. 4E). As expected, the proportion of CD4-CD8-cells in CD3+ cells was reduced following enrichment.

**Figure 4.**
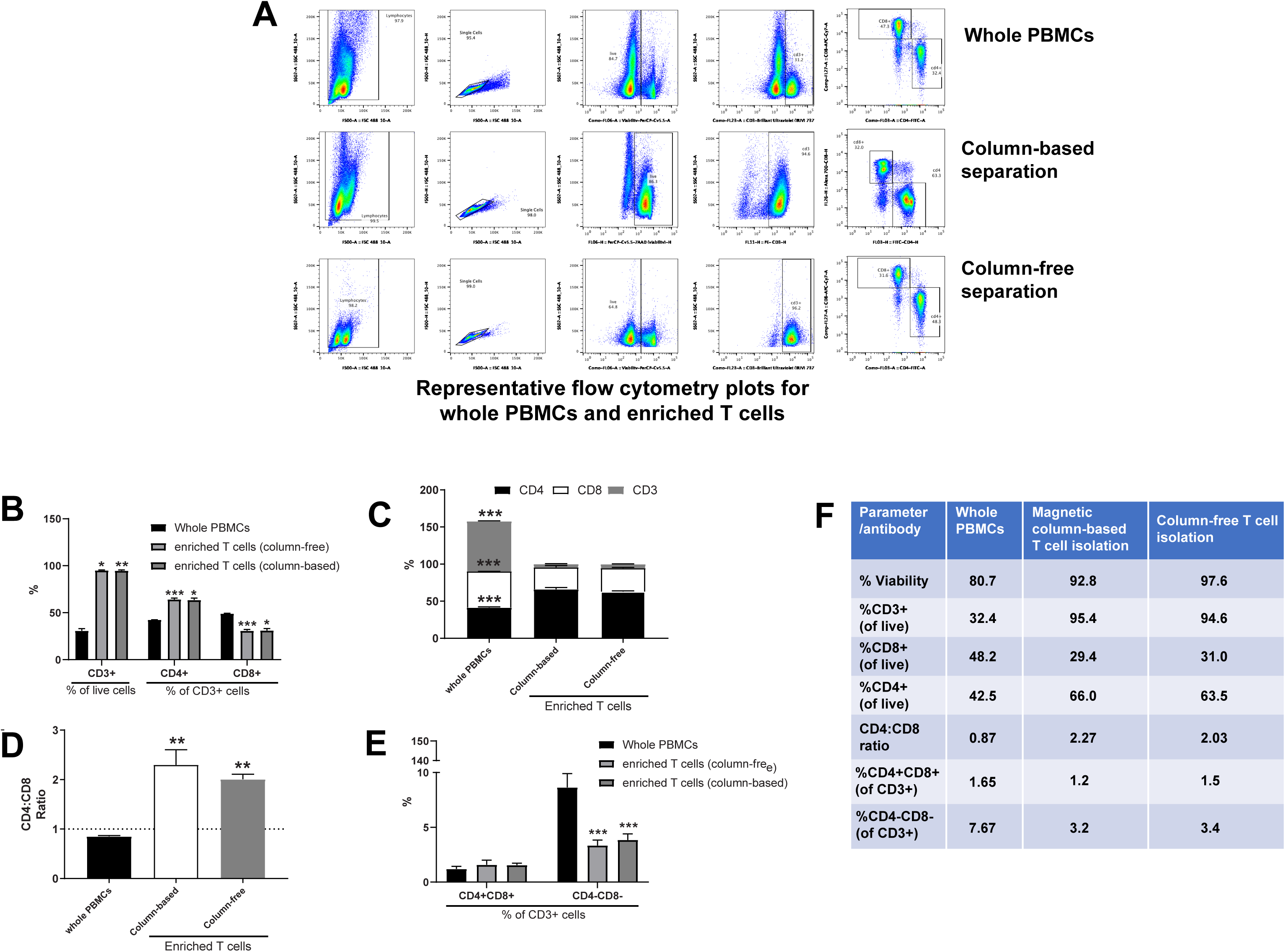
Comparing yield and purity of T cell enrichment methods. T cells were negatively enriched from frozen human PBMCs using either a column-based or column-free magnetic separation method. (A) Following enrichment, whole PBMCs and enriched cells were stained with antibodies for viability and CD3, CD4 and CD8 surface markers. Samples were run through the BioRad ZE2 Cell analyzer. For every sample, gating was as follows: Lymphocyte gate>Singlet gate>Viability gate> CD3+ gate> CD4 and CD8. (B) Bar graph depicting % of CD3+ cells as a percent of all cells, CD4+ cells as a % of CD3+ cells and CD8+ cells as a % of CD3+ cells across all conditions. Two-way ANOVA, with Tukey’s Multiple Comparison Test. 5 *p* < 0.01 and *** *p* < 0.001. n=3. (C) Bar graph depicting % of CD4+, CD8+ and CD3-cells in all cells across all conditions. Two-way ANOVA, with Tukey’s Multiple Comparison Test. *** *p* < 0.001. n=3. (D) CD4:CD8 ratio across all conditions. One-way ANOVA, ** *p* < 0.01, n=3. (E) Comparison of % CD4+CD8+ and % CD4-CD8- of CD3+ cells across all conditions. Two-way ANOVA, with Tukey’s Multiple Comparison Test. *** *p* < 0.001. n=3. (F) Summary table quantifying % viability, % CD3+ cells (of live cells) and % CD4+, CD8+, CD4+CD8+ and CD4-CD8- of CD3+ cells.

In summary, in whole PBMCs, there was an average of 32% CD3+ T cells, of which 48% were CD8+ and 41% were CD4+ cells. On enriching T cells from whole PBMCs, we found that (1) Magnetic-column-based T cell isolation kit resulted in an average of 95% CD3+ T cells, with 29% of those being CD8+ and 66% being CD4+ cells, and (2) Column-free T cell Isolation kit resulted in an average of 94% CD3+ T cells, of which, 31% are CD8+ cells and 63% are CD4+ cells, (3) the ratio of CD4+ to CD8+ cells increased from 0.87 in whole PBMCs to ∼ 2.1 in enriched T cells. And (4) there was no difference in proportion of CD4+CD8+ double positive cells in CD3+ cells across all conditions (Fig. 4F, please refer to Statistical Analyses worksheet for detailed information on means, SD and N). We suggest that care should be taken when analyzing single-cell datasets that derive from pre-enriched T cells as they may have altered numbers of CD4 and CD8 cells compared to whole PBMCs.

#### B cell enrichment

For each human PBMC sample, 2 B cell negative enrichment kits were tested: (1) column-based magnetic separation and (2) column-free magnetic separation. A small aliquot of unmanipulated PBMC sample was run as-is on the flow cytometer alongside enriched cells. For every sample, gates were created for removing doublets (FSC-A vs. FSC-H) and then dead cells using 7-AAD viability stain. Following this, live, single cells were gated on CD19 and CD20 gene expression. Double-positive cells were considered to be B cells (Fig. 5A).

**Figure 5.**
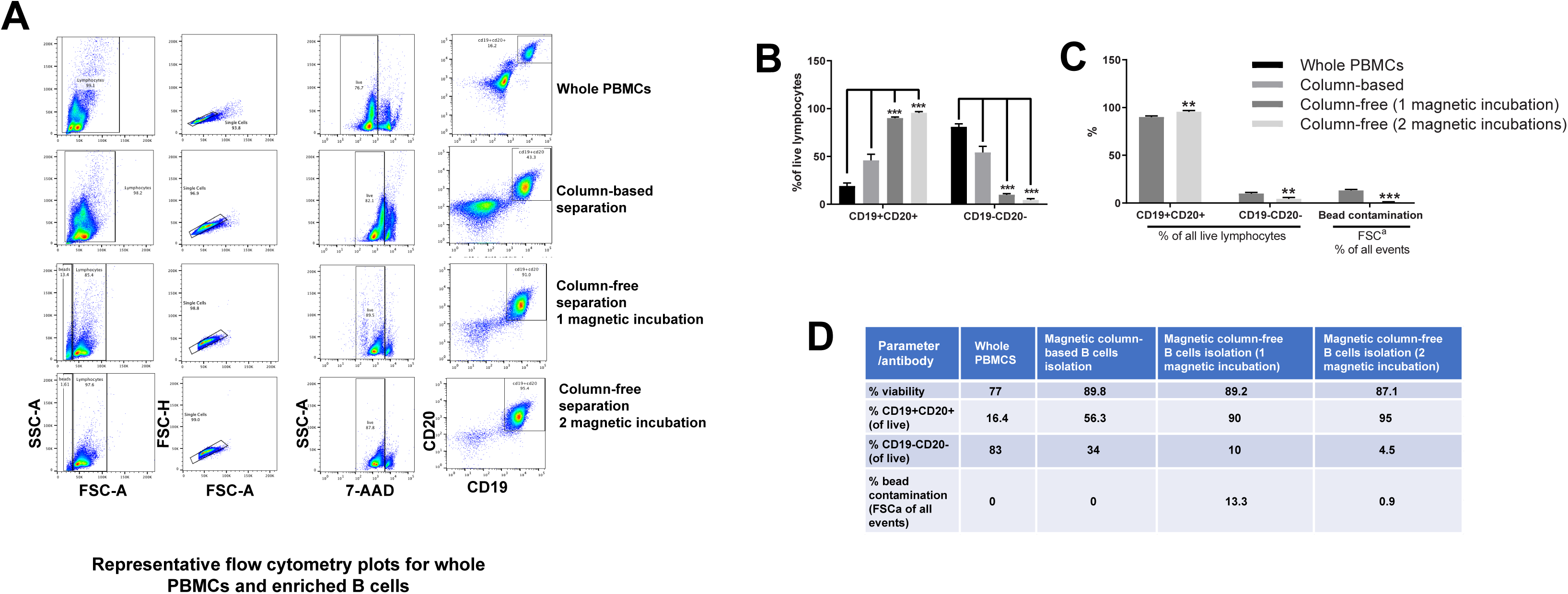
Comparing yield and purity of B cell enrichment techniques. B cells were negatively enriched from frozen human PBMCs using either a column-based or column-free magnetic separation method. (A) Following enrichment, whole PBMCs and enriched cells were stained with antibodies for viability and CD19 and CD20 surface markers. Samples were run through the BioRad ZE2 Cell analyzer. For every sample, gating was as follows: Lymphocyte gate>Singlet gate>Viability gate> CD19, CD20 gate. (B) Bar graph depicting % of CD19+CD20+ and CD19-CD20- cells of all cells across all conditions. (C) Bar graph depicting % CD19+CD20+, CD19-CD20- cells and beads contamination of all cells in 1 magnetic incubation vs. 2 magnetic incubations using Column-free method. (D) Summary table quantifying the % of B cells, non-B cells and beads as obtained from flow analysis across all conditions. Two-way ANOVA, with Tukey’s Multiple Comparison Test. ** *p* < 0.01 and *** *p* < 0.001. n=3.

A small amount of bead contamination was found to be present in all samples that underwent column-free B cell enrichment (Fig. 5A second to last row, red arrow). In order to remedy this, we modified the original protocol and incubated cells with the magnet a second time immediately after 1^st^ incubation and decantation (Fig. 5A, last row).

On comparing the number of B cells across all methods, it was evident that column-free magnetic separation outperformed column-based method, as it significantly increased the yield of B cells and significantly reduced the number of contaminating non-B cells (Fig. 5B, two-way ANOVA, with repeated measures). Additionally, comparing 1 magnetic incubation with 2 magnetic incubations, we found that this not only removed contaminating beads as expected, but it also increased B cell yield significantly and significantly reduced non-B cell number as well (Fig. 5C, two-way ANOVA, with repeated measures).

On quantifying cell numbers, we found that in healthy human PBMCs, with an average viability of 77%, an average of 16% of all cells were B cells, as identified by CD19+CD20+ cells (Fig. 5D, please refer to Statistical Analyses worksheet for detailed information on means, SD and N). All B cell enrichment methods increased the viability to almost 90%. Magnetic-column-based separation resulted in average of 56.3% B cells of all enriched cells. Column-free separation resulted in average of 90% B cells in the enriched population and 13% bead contamination. After incubating with the magnet a second time in the column-free method, B cell purity increased to 95% and bead contamination reduced to 0.9%. The number of contaminating non-B cells were the highest in Column-based method resulted (average of 34%) with Column-free method (1 magnetic incubation) resulting 10% contaminating cells and 2 magnetic incubations resulting in less than 5% contaminating cells.

#### Single Cell analysis on whole PBMCs and enriched B cells

As demonstrated above, the proportion of B cells within frozen human PBMCs is approximately 16%. When performing single cell sequencing, this can translate to 160 cells in 1000 PBMCs or 800 cells in 5000 PBMCs. On the outset, these numbers can seem sufficient for B cell analysis. However, patient-to-patient variability can be very high. Additionally, in cases where sample is stored at less-than-optimum conditions or is obtained from a patient with depleted immune cells due to disease/treatment, getting enough number of B cells might be challenging. As a consequence, obtaining enough cells to represent all subsets in enough numbers can be difficult as well. Hence, we compared the B-cell subtype data (cell abundance, gene expression and repertoire) obtained from single-cell analysis of whole PBMCs versus that of enriched B cells from the same sample. Further, we wanted to test whether 1 vs. 2 magnetic incubations during enrichment process can affect quality of single-cell data at the end. Hence, we performed 5’ gene expression and V(D)J repertoire profiling on whole PBMCs and enriched B cells using the column-free method. We targeted 4000 cells per sample.

Using SPRING single-cell feature reduction and visualization tool [15, 16], we assessed gene expression of markers used to identify different cell subtypes. We used CD19, CD20, CD79A and CD79B as typical markers for B cell identification (Suppl. Fig. 3A), and found that, indeed, there was an enrichment of cells expressing these markers in single-cell data of pre-enriched samples compared to whole PBMCs, as expected.

Similar to our flow analysis data, the effect of incubating cells with magnet twice was clearly seen in the single-cell data of monocyte (CD14) and T cell populations (CD3D, CD4 and CD8A), as they were further depleted in the samples that underwent 2 magnetic incubations (Fig 6A). Next, we separated out B cell clusters from all the samples and visualized them separately on SPRING for the sake of simplification. On deeper investigation of B cell subset markers, we found that the overall proportion of naïve B cells (CD27-, IgD+) increased from 3.5% in whole PBMCs to 57% in enriched B cells (Fig. 6B, 6G), memory B cells (CD27+, IgD-, IgHg2+) increased from 1.4% to 27% (Fig. 6C, 6G) and plasma cells (CD27+, IgD-, CD38+, SDC1+) increased from 1% to 5% (Fig 6D, 6G) of all analyzed cells. Immunoglobulin heavy chain genes IGHG2 and IGHA1 genes were also enriched in the plasma cell population in the enriched B cells (Fig. 6E). Further, IGHE gene expression that was present in the memory cells of enriched B cells, was not detectable in the B cells from whole PBMCs. Similarly, IGHJ genes (1-3) were completely undetectable in the whole PBMC B cells. Finally, we saw a cell cluster expressing IGLL1, that was not present in whole PBMCs (Fig. 5F). This cluster was also enriched for the hematopoietic stem cell marker, CD34 gene expression. Thus, hematopoietic stem cells were only detectable in enriched B cells, and not whole PBMCs, likely due to their relatively small proportion in PBMCs. In summary, pre-enriching B cells from whole PBMCs drastically increased the proportion of B cells analyzed from 6% to 89% and reduced the proportion of T cells and monocytes from 62% to 1% and 27 % to 0.1% respectively of all cells (Fig. 6G). Furthermore, B cell subtypes such as naïve cells, memory B cells and plasma cells were also increased in proportion in enriched B cell samples. CD34+ cells, which were not detectable in whole PBMCs, were visible in enriched B cells (0% vs. 0.34%, not shown in table).

**Figure 6.**
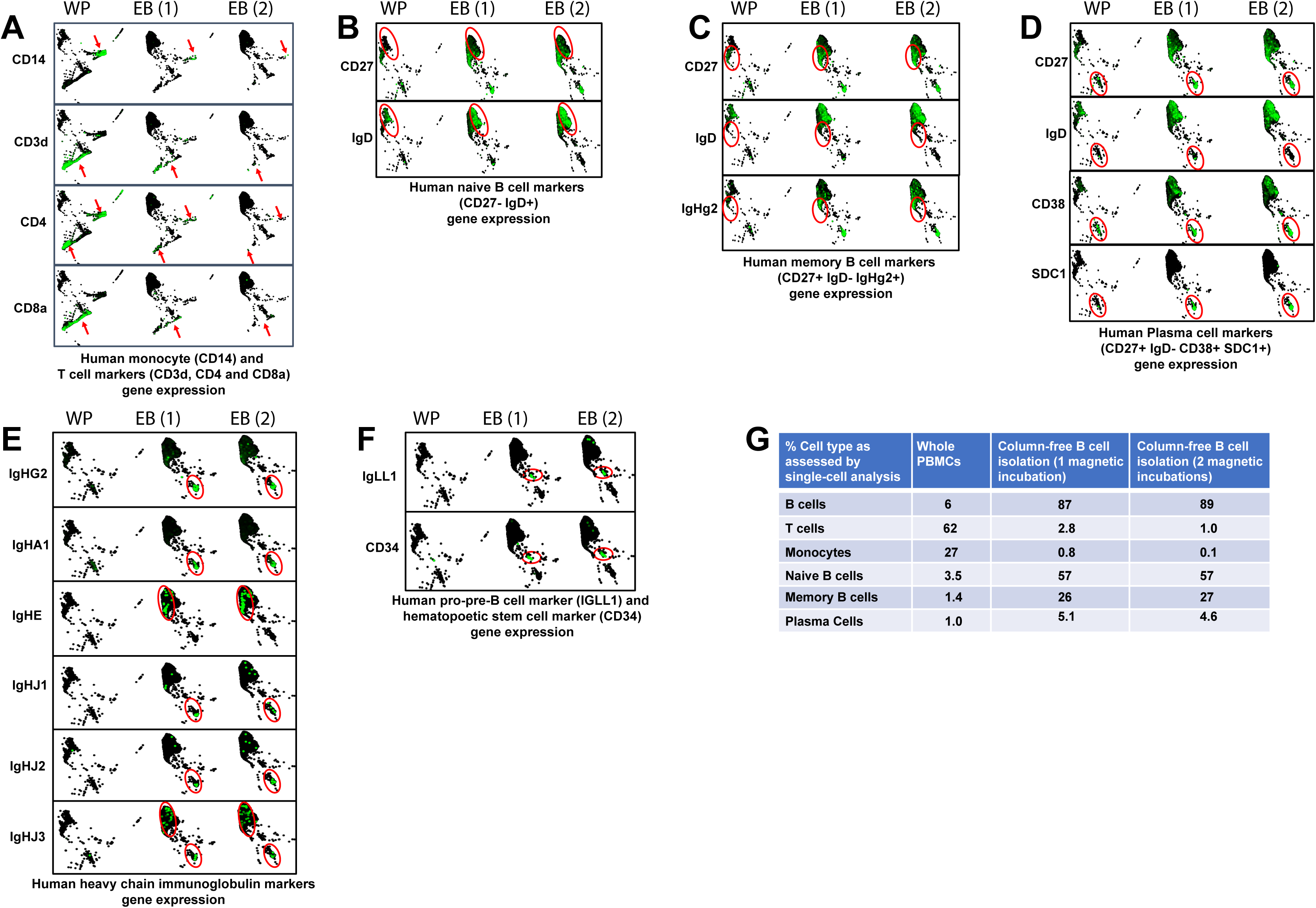
Assessing single-cell gene expression for whole PBMCs, enriched B cells (1 and 2 magnetic incubations). Single-cell Visualization Plots for whole PBMCs, enriched B cells (1 magnetic incubation or 2 magnetic incubations) on SPRING. (A) Commonly used genes as markers for identification of monocytes and T cells: CD14, CD3D, CD4, CD8A. Arrows depict specific areas of depletion in enriched cells in 1 and 2 magnetic incubations. (B) Gene expression of CD27 and IgD to identify naïve B cells (CD27-IgD+) in red circle. (C) Gene expression of CD27, IgD and IgHg2 to identify memory B cells (CD27+IgD-IgHg2+) in red circle. (D) Gene expression of CD27, IgD, CD38 and SDC1 to identify plasma cells (CD27+, IgD-, CD38+, SDC1+) in red circle. (E) Gene expression for heavy chain immunoglobulin markers. IgHg2 and IgHA1 are expressed in plasma cells and IgHE is expressed in memory cells of enriched B cells. IgHJ genes are detectable in enriched B cells, but not in whole PBMCs. (F) IgLL1 and CD34 gene expression as a marker for hematopoietic stem cells are visible in enriched B cells, and not in human PBMCs. (G) Summary table quantifying the % of different cell subtypes in single-cell dataset of whole PBMCs and enriched B cells (1 and 2 magnetic incubations) as analyzed in SPRING.

Next, we assessed different BCR clonotypes expressed using V(D)J repertoire profiling. Using an in-house cell type annotation tool, we identified major cell subtypes in whole PBMC and enriched B cell samples on SPRING (Fig. 7A, top). Next, we removed any cells that did not express any BCR clonotypes from the analysis (Fig. 7A-bottom). What remained were cells expressing rare clonotypes (light grey dots), and some commonly expressed clonotypes (multi-colored dots). Interestingly, on quantifying the percentage of cells expressing any BCR clonotype across different cell subsets, we found that 99% of all BCR clonotypes were expressed specifically on B cells or plasma cells in the enriched B cell population (Fig. 7B). However, in whole PBMCs, approximately 25% of all BCR clonotypes were expressed nonspecifically on non-B cells such as T cells or monocytes. On the contrary, on looking at low frequency clonotypes, we found that most of the rare clonotypes were expressed on B cells.

**Figure 7.**
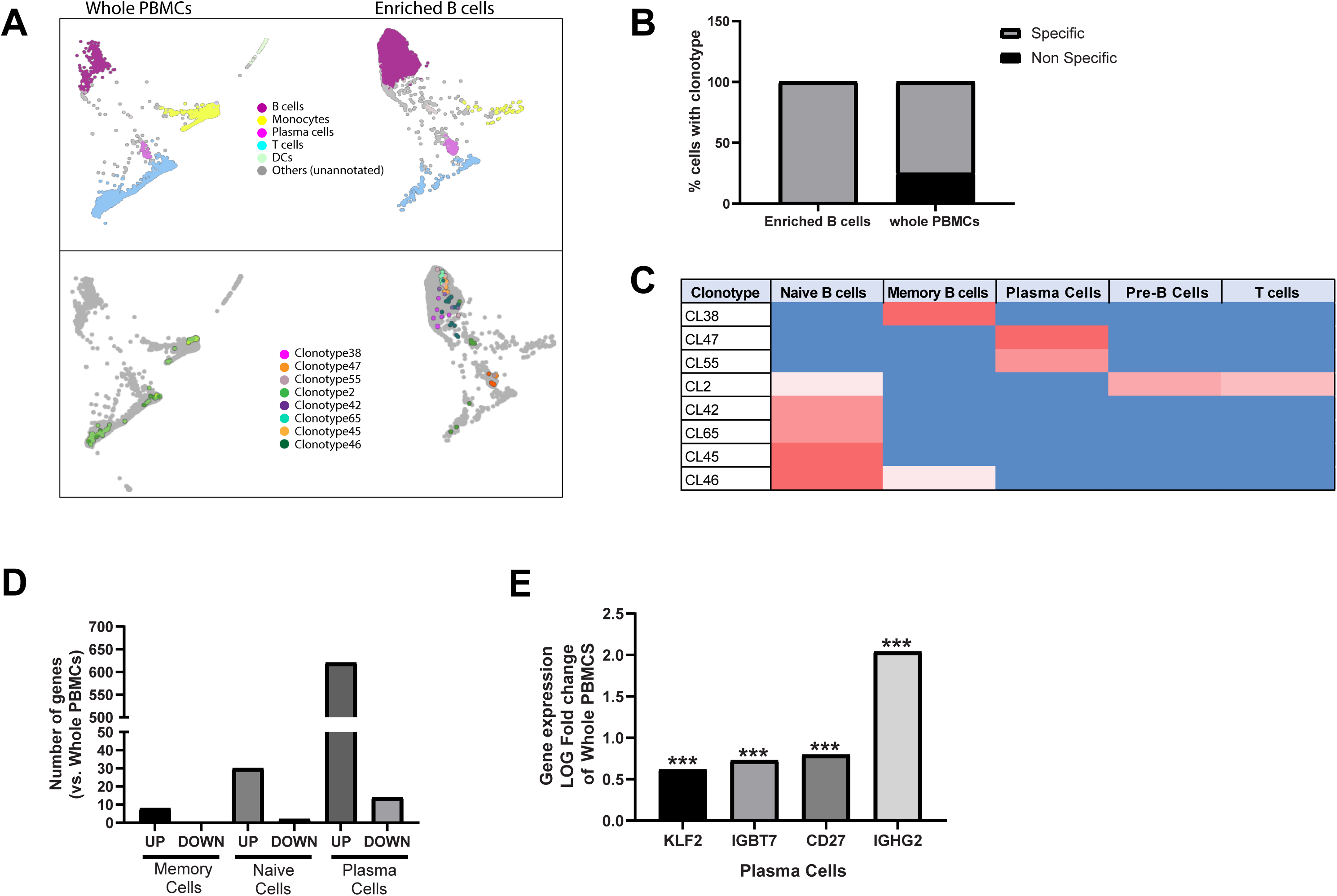
Single-cell BCR repertoire profiling of whole PBMCs, enriched B cells (1 or 2 magnetic incubations). Single-cell V(D)J repertoire profiling of whole PBMCs, enriched B cells (1 magnetic incubation or 2 magnetic incubations) on SPRING. For whole PBMCs, B cell V(D)J target enrichment was performed from the cDNA after which library was prepared. (A, top) Major cell subtypes highlighted on whole PBMCs and enriched B cells using an in-house cell-type annotation tool on SPRING. (A, bottom) BCR clonotypes distribution across whole PBMCs and enriched B cells. (B) Quantification of % of cells expressing BCRs specifically in B cells and non-specifically in non-B cells. (C) Summary of distribution of major BCR clonotypes across B cell subtypes in enriched B cell samples. (D) Number of genes with significantly different log Fold change in memory B cells, naïve B cells or plasma cells of enriched population compared with those of whole PBMCs. (E) Log fold gene expression change of select genes that are significantly upregulated in plasma cells of enriched cells compared with those of whole PBMCS. Wilcoxon Rank Sum Test, adjusted p-value is based on multiple test correction using the Bonferri method.

This was true for both whole PBMCs and enriched B cells (Suppl. Fig. 3C). The “missing” clonotypes were mostly in non-B cells such as T cells or monocytes. This suggested that for whole PBMCs library preparation, the V(D)J target enrichment might be nonspecific and result in some spurious expression of high-frequency BCR clonotypes on non-B cells.

Next, we analyzed in detail the cell-subtype expression of the BCR clonotypes that were highly expressed in enriched B cells (Fig. 7C). One of the individuals profiled exhibited CL38 uniquely in memory B cells. All other frequent clonotypes were detected in the other individual. Naïve B cells had the highest diversity of BCR expression, with four unique clonotypes expressed (CL42, CL65, CL56 and CL45), whereas plasma cells uniquely expressed three BCRs: CL47, CL55 and CL7. We found that clonotype CL2 was non-specifically expressed in naïve B cells and T cells, while CL46 was expressed in both naïve B cells and memory B cells. None of the highly expressed clonotypes in enriched B cells were present in B cells of whole PBMCs. Thus, we found that performing V(D)J target enrichment on whole PBMCs can result in frequent clonotype information which would not necessarily be obtained from pre-enriched B cells. Finally, we assessed if there were any significantly differentially expressed genes in memory cells, naïve cells and plasma cells of enriched B cells vs. those of whole PBMCs (Fig. 7D). There were less than 8 genes that were significantly up-regulated and no genes significantly down-regulated in memory cells of enriched B cells compared to those of whole PBMCs. There were 30 genes significantly upregulated and only 2 genes significantly downregulated in the naïve cells of enriched cells compared to those of whole PBMCs. However, in when looking at plasma cell gene expression, we found 620 genes significantly up-regulated, and 14 genes significantly down-regulated in enriched cells as opposed to those in whole PBMCs. This suggested that pre-enrichment does not affect the gene expression profile of memory and naive B cells significantly compared to whole PBMCs. However, pre-enrichment does affect the expression of plasma cell genes. This could be likely due to increased number of cells being detected overall in the enriched population compared to the relatively low percentage of plasma cells found in whole PBMCs. Particularly, we found important functional genes to be upregulated significantly in the plasma cells of enriched population compared to those in whole PBMCs, such as KLF2, IGBT7, CD27 and IGHG2 (Fig. 7E). For full list of differentially expressed genes in memory B cells, naïve B cells and plasma cells across whole PBMCs and enriched B cells, as well as detailed statistical analyses, please refer to “Supplementary_DEG” excel worksheet.

## A quantitative worksheet

To sum up our observations, we developed a calculation aid which allows researchers to explore cell requirements and yields at different workflow steps (Supplemental worksheet: <u>Calculator</u>). Based on our data presented above (Figures 4-8), we calculated the percentage of cells lost at every step of single-cell workflow, including washes, sample-clean-up and pre-enrichment for T or B cells. Furthermore, we also predict number of cells lost in case of consecutive steps of sample clean-up followed by pre-enrichment. Following thawing and washing of PBMCs, user is encouraged to count cells manually using hemacytometer. For workflo multiple samples are available, AO/PI based automated counters may be used. Following this, user can input their TOTAL cell number in the worksheet, in the orange box. The worksheet will automatically populate to show potential number of cells that would remain following sample clean-up procedures, T or B cell enrichments, or sample cleanup + T or B cell enrichments. The graph depicts the cell yield post-cleanup for different methods at different starting cell numbers (X-axis). As the clean-up performance, i.e. increase in % viability varies at different starting % viability values, this cannot be predicted accurately in the calculator.

## Discussion

For a successful single-cell workflow, it is crucial to know the cell count and viability of the sample accurately prior to single-cell encapsulation. Trypan blue (TB) is a commonly used dye to distinguish between live and dead cells. One of the most common methods of counting cells using TB is manual counting using hemacytometer slide. As this method involves counting of cells by a researcher under a microscope, this is a time-consuming process and may introduce operator-dependent variability. Further, red blood cell (RBC) contamination can make counting process all the more challenging. There are several automated counters that use the same TB dye but remove the human error and can save time. Furthermore, there are features in the software that reduce counting errors introduced by RBC contamination by defining a cell size range for counting. To test whether automated counters that use TB as a dye are reliable, we compared PBMC cell counts from 2 automated counters against that of manual counting. We found that both automated counters under-estimated dead cells, but over-estimated live cells. Hence, the reported viability values were inflated compared to manual counting. This poses as a dangerous risk, because if a reported value is 90%, when in reality it is only 65%, it can reduce the quality of single-cell data drastically. Hence, we recommend avoiding the use of trypan-blue based automated counters for single-cell workflows involving PBMCs. Intriguingly, since TB automated counters are used as the basis for the 90% viability cutoff recommended in the popular 10X Genomics® single-cell protocol, it is possible that the true viability requirements are not as stringent. Conversely, the 90-95% recommendation in the 1CellBio protocol is based on hemacytometer and may be unnecessarily high.

In spite of the obvious advantages of manual counting as described above, this method can be tedious and time-consuming when many samples are to be counted. For this reason, we investigated whether fluorescence-based automated counters would provide a more accurate and reliable cell count compared to TB-based automated methods. Fluorescence-based counting is generally considered more accurate because dead and live cells can be stained differentially using a combination of dyes that fluoresce at different wavelengths. For example, when Acridine Orange and Propidium Iodide (AO/PI) are used for staining cells, AO permeates both live and dead nucleated cells, whereas PI enters only dead nucleated cells. As a result, all live nucleated cells fluoresce green and all dead nucleated cells fluoresce red, due to a phenomenon called as Forster resonance energy transfer or FRET [20]. RBCs are specifically not stained by AO/PI due to their lack of nucleus[21]. We compared PBMC cell counts from two automated fluorescence-based counters with those of manual TB-based counting. We found that both fluorescence-based automated counters reported viability values comparable to that of manual counting. We found an overall significant difference in live cells per ml counts across different methods, but this did not translate to differences in total cells per ml or % viability. These results are consistent with an earlier study which compared trypan blue vs. AO/PI for counting of hematopoietic progenitor cells (HPCs) from the bone marrow in the context of bone marrow transfusion in the clinical setting [22]. In that study, AO/PI-based cell counting measurements had a stronger correlation with predicted viability values as well as with CFU-GM concentration (progenitor cell function) of HPCs compared to TB. As AO/PI displayed better stability and less over time compared to TB, authors concluded it to be the more reliable assay. We therefore recommend researchers to use fluorescence based automated counters when sample throughput is a concern. Manual counting is the most reliable method when only a few samples are being handled.

Once the viability of a cell suspension is established, the researcher has to make a decision on whether to move forward with single-cell workflow or apply dead-cell removal method to clean-up the sample. As already mentioned, viability of 90% or more is often cited as a cutoff for proceeding with single-cell microfluidic workflows. Furthermore, any improvement in viability will result in better quality data due to reduced background contamination. Hence, we wanted to investigate which is the best method to remove dead cells from a sample with poor viability and potentially low quantity. We found that magnetic-column-based bead separation resulted in the highest viability post-clean-up, whereas washes low-speed spin (as recommended by 10X Genomics® protocol) result in least clean-up efficiency. Conversely, washes with low-speed spins are most ideal when starting with a low amount of sample as it results in the least amount of cell loss. All other methods, and specifically the magnetic beads kits, drastically reduce the total number of cells. Hence, we recommend using these only if viability is a significant concern and if ample starting material is available, with the caveat that there is severe loss of total cells at the end (Supplementary <u>Calculator</u> Worksheet). Otherwise, three washes with low-speed spins might be sufficient to bring the viability up to a reasonable level for single-cell workflows. As a general guideline, we recommend performing dead cell removal using column-based magnetic kit <u>only</u> if starting material is more than ∼2.5 million total cells. Thus, enough cells will be available for subsequent lymphocyte pre-enrichment, if necessary. Specifically, approximately 100,000 B cells may be expected from 2.5 million starting PBMCS, following column-based dead cell removal and pre-enrichment (See Calculator worksheet). It is also of utmost importance to resuspend cell pellets between washes gently with wide-bore pipette tips to minimize cell death due to attrition.

The concentration of lymphocytes within frozen human PBMCs is approximately 45%. Of this, approximately 10-15% are B cells and remaining are T cells. However, these numbers may vary across individuals. Additionally, in case of immunological diseases, the number of lymphocytes could potentially be affected, such as in cases of cancer or HIV. Certain treatments can also deplete lymphocytes further. In the context of single-cell sequencing where in only a few thousand cells are analyzed, the number of lymphocytes could be down to only a few tens per sample. Pre-enrichment might be necessary to get enough cells for analysis, or desirable in addition to PBMC profiling given enough starting material (Supplementary Calculator Worksheet). By enriching these cells, one can (1) get more information about their subtypes, (2) potentially clean-up low viability samples, and (3) obtain accurate information about repertoires on a single-cell basis. Hence, we sought to determine the best method to enrich for these cells from a previously frozen human PBMC sample. For both cell types we used negative enrichment kits, so as to maximally preserve the heterogeneity of the target cell type and avoid biasing the selected population towards high levels of specific positive markers. Although the risk for contaminating cells being collected is high following negative enrichment, this method leaves the cells relatively untouched compared to positive enrichment separation [23].

For T cell enrichment, the performance of both column-based and column-free kits were comparable. There was also no evidence of bead contamination in the enriched cell population following the use of the column-free kit. However, an interesting observation was the inversion of CD4 to CD8 cell ratio in the enriched populations of both the T cell enrichment kits. The ratio of CD4:CD8 T cells within CD3+ population in whole PBMCs was found to be <1. However, this ratio increased to almost 2 in the enriched CD3+ T cells. We therefore generally recommend against enrichment of T cells from whole PBMCs by these methods. In reality, enrichment of T-cells is rarely necessary as they constitute almost half of the PBMC cell population. However, disease condition and treatments might lower T-cell abundance, in which case these or alternative enrichment solutions may be sought, depending on the specific scientific objective.

We found that for B cells, column-free magnetic separation provided the best yield and purity. We modified the protocol in order to get rid of contaminating beads. Single-cell analysis showed that not only did this modification increase the yield of B cells, but it also reduced contaminating cells such as monocytes and T cells. Furthermore, on comparing immunoglobulin expression across whole PBMCs and enriched B cells it was evident that certain heavy chain immunoglobulin genes (IGHG2. IGHA, IGHE and IGHJ) were not detected in whole PBMCs, which were detected in enriched B cells. Especially, expression of immunoglobulin genes in plasma cells was drastically reduced in whole PBMCs. This could be simply because not enough cells were sampled. Similarly, we detected the presence of hematopoietic stem cells through the marker CD34 gene expression [24]. The difference between BCR clonotype expression in whole PBMCs and enriched B cells was striking. Not only were the number of clonotypes detected higher in enriched cells, but the specificity was also much improved. The detection of BCR clonotypes in T cells of whole PBMCs can be a cause for concern. Finally, we found only a few genes that were significantly changed in the memory cells and naïve cells, and hence we conclude that pre-enrichment does not significantly affect the gene expression patterns of these cells. However, 634 genes were significantly changed in plasma cells of enriched cells, compared to those in whole PBMCs. Of these, several genes important for plasma cell function, activation and homing were upregulated. For example, KLF2 and ITGB7 are important for their function in plasma cell homing [25, 26], whereas CD27 is important for plasma cell activation [27]. We believe that because the relative abundance of plasma cells is much higher in enriched cells, it could result in richer gene expression data.

Finally, we present a table (Table 1) to summarize our findings in a concise way, as well as provide recommendations for best practices for each step in the workflow. Specifically, we recommend using a manual hemacytometer wherever possible for obtaining highly accurate cell counts and viability measurements. We also propose multiple ways for sample clean-up for poor viability samples, based on the amount of starting material and initial viability post-thawing. We also find that pre-enrichment of B cells provides most information on B cell expression, cell subtype and VDJ repertoire compared to whole PBMCs. Finally, using T cell enrichment kits comes with caveats because of the altered ratio of CD4:CD8 cells compared to that in whole PBMCs. Overall, our work provides specific guidelines for processing of clinical samples for single-cell analysis, and a quantitative aid for determining the possible workflows given the amount of starting material. We also believe these data will help scientific research community at large due its applicability across different areas of immunology.

**Table 1.**

| Purpose | Methods used | Result | Recommendations |
| --- | --- | --- | --- |
| **Viability assessment** | 1. Manual Hemacytometer<br>2. TB-based automated counters<br>3. AO/PI-based automated counters | Manual hemacytomter counts are most reliable.<br>TB-based automated counters over-estimate viability.<br>AO/PI-based automated counts are comparable to manual counting. | In case of few a samples, manual counting is recommended.<br>For counting of multiple samples, AO/PI-based automated coutners may be used. |
| **Sample clean-up** | 1. Low-speed spin washes<br>2. Column-based magnetic clean-up<br>3. Column-free magnetic clean-up<br>4. Density gradient based clean-up | Low speed-spin washes are least effective in improving viability, but result in the least amount of cell loss.<br>All kit-based dead-cell-removal methods result in significant cell loss (~50%), however, column-based method is most effective in improving viability. | When starting with low total cell numbers ($< 2.5 \times 10^6$), low-speed spin-washes may be used. If more than $2.5 \times 10^6$ cells are available, column-based clean-up methods may be used so as to result in enough cells for subsequent lymphocyte enrichment, if necessary. |
| **T cell negative enrichment** | 1. Column-based magnetic separation<br>2. Column-free magnetic separation | Both column-based and column-free negative enrichment kits perform comparably resulting ~95% T cells following enrichment. However, the CD4:CD8 ratio increases from $<1$ to $>2$ following enrichment. | Avoid T cell pre-enrichment from whole PBMCs, unless working with special case samples (such as PBMCs with depleted T cells due to disease or treatment). |
| **B cell negative enrichment** | 1. Column-based magnetic separation<br>2. Column-free magnetic separation | Column-based negative enrichment results in only 56% B cells , whereas column-free negative enrichment results in $>95\%$ B cells in enriched population. | Column-free magentic separarion, with two magnetic incubations, is recommended for obtaining high purity B cell enrichment. |
| **5' gene expression and BCR profiling of whole PBMCs vs. enriched B cells** | 1. 5' gene expression, BCR target enrichment and BCR profiling of whole PBMCs.<br>2. 5' gene exprression and BCR profiling of enriched B cells | Heavy-chain immunoglobulin genes are not detectable in plasma cells of whole PBMCs.<br>Non-specific clonotype expression is observed in whole PBMCs, but not in enriched B cells.<br>634 genes significantly changed in plasma cells of enriched B cells compared to those in whole PBMCs. | We recommend pre-enriching B cells from whole PBMCs prior to single-cell analysis, to obtain most accurate information on BCR clonotypes as well as gene expression from B cells and their subtypes. |

## Supporting information

Calculator

Statisitcal_analyses

DEG_analyses

## Acknowledgements

The authors would like to thank Dr. Li Li at Sanofi for kindly providing all frozen human PBMCs samples used in this project. Special thanks to Dr. Alexei Protopopov and his team: Dr. Emma Wang and Maximilian Rogers-Grazado in the Sanofi Oncology-Genomics Department for their help with sequencing single-cell transcriptomics libraries.

---

**Supplementary Figure 1:**
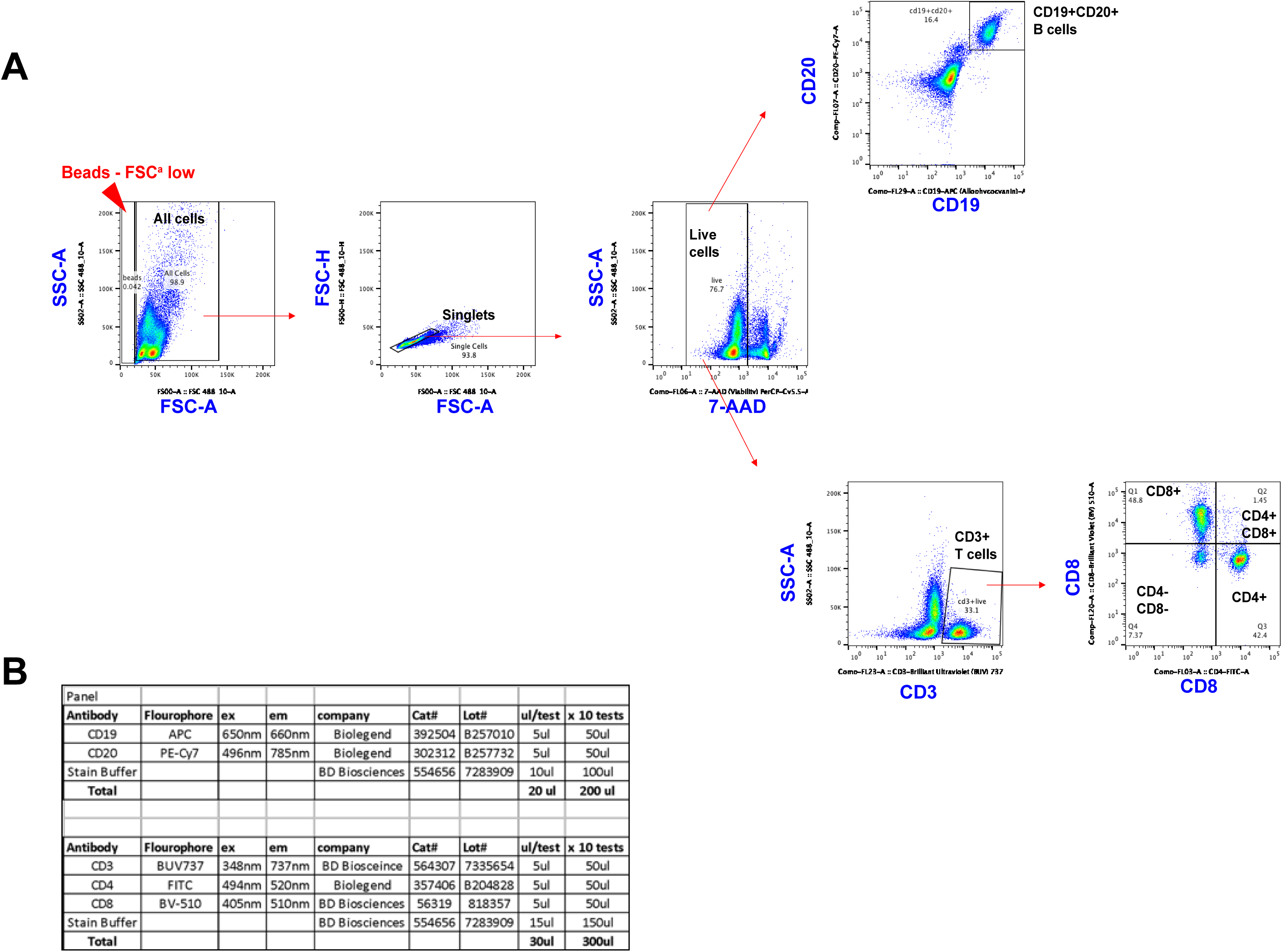
All flow analysis was performed on FlowJo software using a linear scale. All cells were first collected by drawing a gate across forward scatter area (FSC-A) and all side scatter area (SSC-A). Care was taken to avoid the very low FSC-A events on the left. This area mainly consisted of any contaminating beads from enrichment of cells and were identified by the “no or low FSC-A” with a wide SSC-A spread. Singlets were identified from <u>all cells</u> by FSC-A by FSC-H (height). Any cells that were low on FSC-H but high on FSC-A were excluded as doublets. Following this, viability was assessed by 7-AAD on X-axis and SSC-A on the Y-axis. Cells low in 7-AAD expression (less than 10^3^) were considered to be viable and labeled as “live cells”. From the live cells, (1) B cells were identified by plotting CD19 vs. CD20, and gating only on the double positive cells, i.e. cells high in both CD19 and CD20 expression, and (2) T cells were identified by CD3 expression on X-axis and SSC-A on the Y axis. Only cells high for CD3+ expression (more than 10^3^) were labeled as T cells. From CD3+ gate, CD4 vs. CD8 were plotted on X and Y axis respectively and a “Quad” gate was drawn resulting in the following quadrants: (a) CD4-CD8+, (b) CD4+CD8+, (c) CD4+CD8- and (d) CD4-CD8-. For every sample, gates were first drawn on whole PBMCs, and then copy-pasted onto enriched cells for the same sample. Efforts were taken not to modify gates post pasting. See Supplementary Figure 1 for a typical gating strategy example.

**Supplementary Figure 2.**
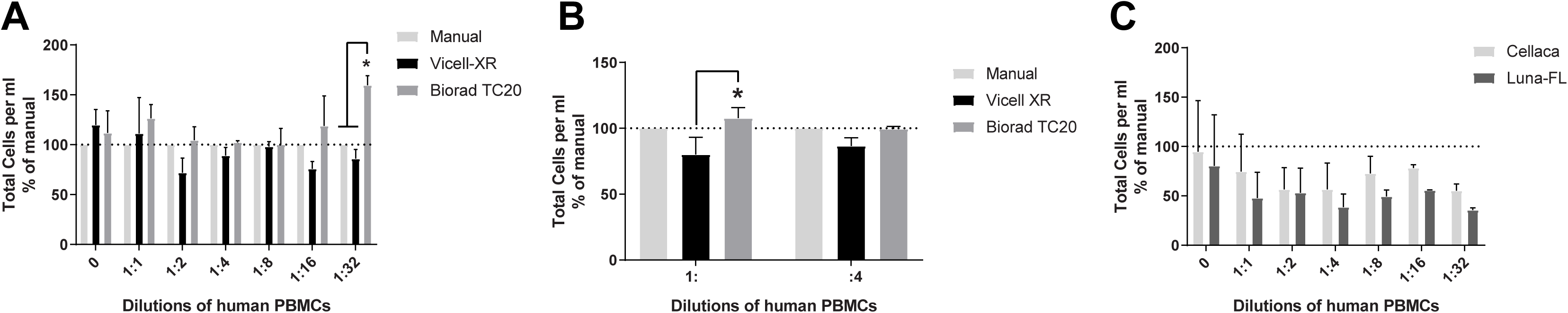
Human PBMCs were diluted using 0.04% BSA/PBS solution and each dilution was counted using Manual hemacytometer, automated TB-based counters or automated AO/PI-based counters. (A) total cells per ml counted by manual hemacytometer vs. TB-based automated counters across different serial dilutions, reported as % of manual counts. Two-way ANOVA, with matched samples, followed by Tukey’s Multiple Comparison test (n=3) (B) total cells per ml counted by manual hemacytometer vs. TB-based automated counters across 2 dilutions 1:2 and 1:4, reported as % of manual counts. Two-way ANOVA, with matched samples, followed by Tukey’s Multiple Comparison test (n=6) and (C) total cells per ml for manual counting vs. AO/PI-based automated counters, reported as % of manual counts. Two-way ANOVA, matched samples followed by Tukey’s multiple comparison test (n=4). For detailed statistics refer to Supplementary information.

**Supplementary Figure 3:**
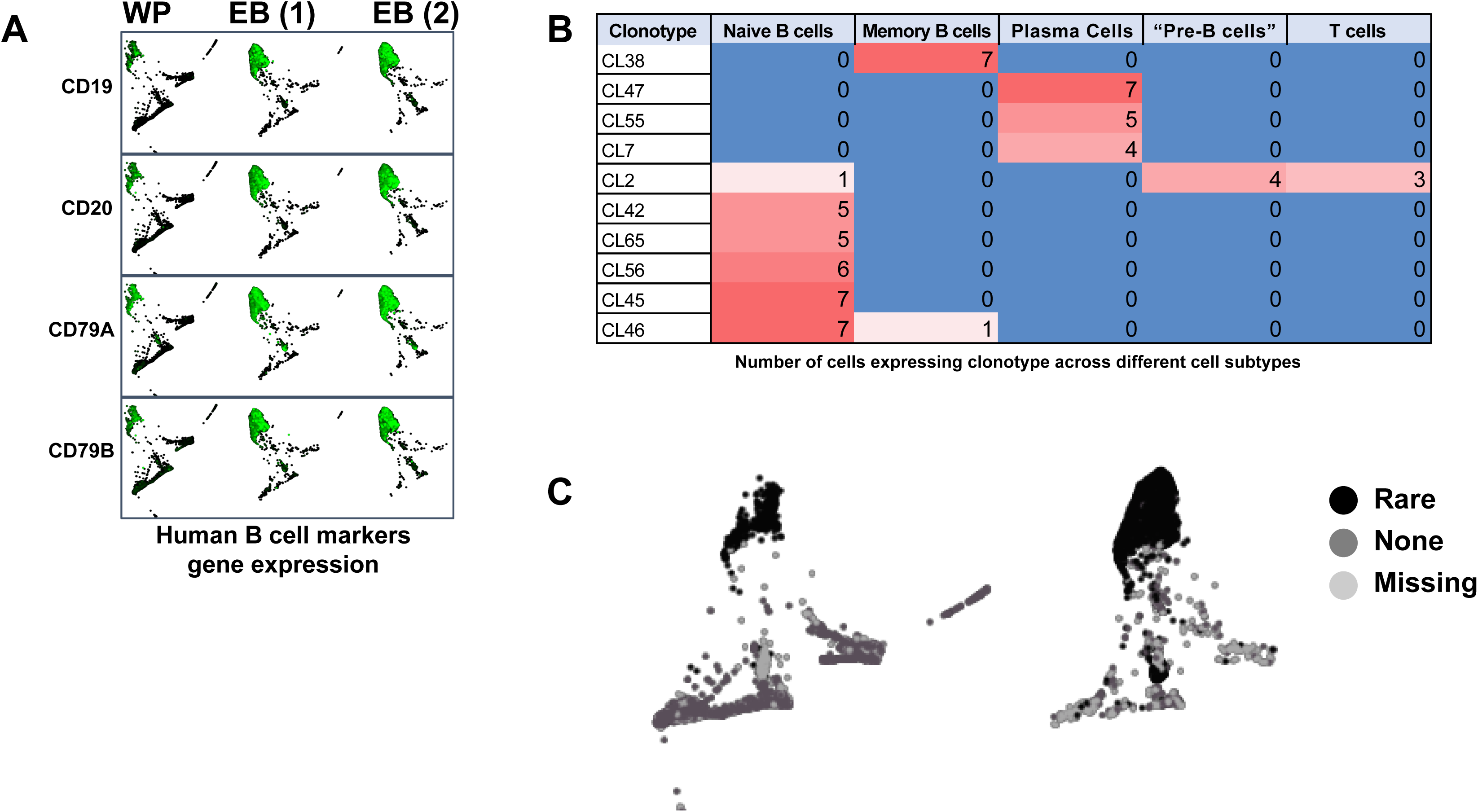
(A) Single-cell Visualization Plots for whole PBMCs, enriched B cells (1 magnetic incubation) and enriched B cells (2 magnetic incubations) on SPRING. Commonly used genes as markers for identification of B cells: CD19, CD20, CD79A and CD79B. (B) Single-cell V(D)J repertoire profiling of whole PBMCs, enriched B cells (1 magnetic incubation) and enriched B cells (2 magnetic incubations). Table depicts number of cells in each cell subtype expressing specific BCR clonotypes.

